# Frequency switching between oscillatory homeostats and the regulation of p53

**DOI:** 10.1101/2019.12.31.891622

**Authors:** Peter Ruoff, Nobuaki Nishiyama

## Abstract

Homeostasis is an essential concept to understand the stability of organisms and their adaptive behaviors when coping with external and internal assaults. Many hormones that take part in homeostatic control come in antagonistic pairs, such as glucagon and insulin reflecting the inflow and outflow compensatory mechanisms to control a certain internal variable, such as blood sugar levels. By including negative feedback loops homeostatic controllers can exhibit oscillations with characteristic frequencies. In this paper we demonstrate the associated frequency changes in homeostatic systems when individual controllers in a set of interlocked feedback loops gain control in response to environmental changes. Taking p53 as an example, we show how the Per2, ATM and Mdm2 feedback loops -interlocked with p53-gain individual control in dependence to DNA damage and how each of these controllers provide certain functionalities in their regulation of p53. In unstressed cells, the circadian regulator Per2 ensures a basic p53 level to allow its rapid up-regulation in case of DNA damage. When DNA damage occurs the ATM controller increases the level of p53 and defends it towards uncontrolled degradation, which despite DNA damage, would drive p53 to lower values and p53 dysfunction. Mdm2 on its side keeps p53 at a maximum level to avoid premature apoptosis. However, with on-going DNA damage the Mdm2 set-point is increased by HSP90 and other p53 stabilizers leading finally to apoptosis. An essential aspect in p53 regulation at occurring cell stress is the coordinated inhibition of ubiquitin-independent and ubiquitin-dependent degradation reactions and the increasing stabilizing mechanisms of p53. Whether oscillations serve a function or are merely a by-product of the controllers are discussed in view of the finding that homeostatic control of p53, as indicated above, does in principle not require oscillatory homeostats.

## Introduction

The concept of homeostasis is central to our understanding how organisms and cells adapt to their environments and thereby maintain their stability [1–3]. With the development of cybernetics [4,5] control engineering concepts were, for the first time, applied to biological systems [6,7]. With the advancement of molecular biology, robust control theoretic methods were applied at the molecular level, such as integral reign control [8], alongside with integral feedback [9–11], and systems biology methods [12,13]. It became clear that certain reaction kinetic conditions are necessary for the occurrence of integral control leading to the robustness of the feedback controller. These conditions include zero-order kinetics [9,10,14–19], autocatalysis [20–22], and second-order (bimolecular/antithetic) reaction [23,24], which were implemented into various controller motifs, synthetic gene networks, and other negative feedback structures [18,25–27].

A particular interesting aspect is that, under certain conditions, the homeostatic controllers may become oscillatory and preserve, if integral control is present, their homeostatic property by keeping the *average value* of the controlled variable at its set-point [28]. While the occurrence of oscillations is generally avoided in control engineering, natural systems show generally oscillatory behaviors, as found in circadian or ultradian rhythms [29–31].

In this paper we show how a set of inter-connected negative feedback loops maintain robust homeostasis in a controlled variable both under non-oscillatory and oscillatory conditions. We show that each controller has its own frequency and how frequency switching between controllers occur dependent on the perturbation level of the controlled variable. We then show how combined motif 3, motif 1, and motif 5 type of controllers [18] reflect aspects of p53 regulation by Per2, ATM, and Mdm2, respectively. Dependent whether p53 degradation or synthesis is dominant, and dependent whether controllers are oscillatory, either low-level p53 circadian rhythms or higher-level p53 ultradian oscillations can be observed.

## Materials and methods

Computations were performed by using the Fortran subroutine LSODE [32]. Plots were generated with gnuplot (www.gnuplot.info) and edited with Adobe Illustrator (adobe.com). To make notations simpler, concentrations of compounds are denoted by compound names without square brackets. Time derivatives are generally indicated by the ‘dot’ notation. Concentrations and rate parameter values are generally given in arbitrary units (au). In the Supporting Information a set of MATLAB (S1 Matlab), gnuplot, and avi-files (S1 & S2 Gnuplot) are provided to illustrate modeling results.

## Results and discussion

### Cannon’s definition of homeostasis and its realization by inflow and outflow controllers. The nonoscillatory case

Cannon defined homeostasis as the result of coordinated physiological processes, which maintain most of the steady states in organisms by keeping them within narrow limits [33]. One of the typical examples are human blood calcium levels, which, throughout our lifetimes are kept between approximately 9 to 10 mg Ca per dl blood. When levels are outside that range serious illness or death may occur.

The combination of inflow and outflow controllers having integral control [18] allows to keep a regulated variable within such strict limits. Fig 1 shows the arrangement between two collaborative controllers where the set-point of the inflow controller 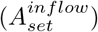 ensures for the lowest tolerable concentration of the controlled variable *A*, while the outflow controller does not allow that *A*-levels exceed the outflow controller’s set-point 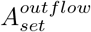. It should be pointed out that not all inflow/outflow controller combinations [18] will lead to a set of collaborative controller pairs, because set-point values and the individual controllers’ on/off characteristics need to match; otherwise the controllers may work against each other and integral windup may be encountered [18], as described in more detail below.

**Fig 1.**
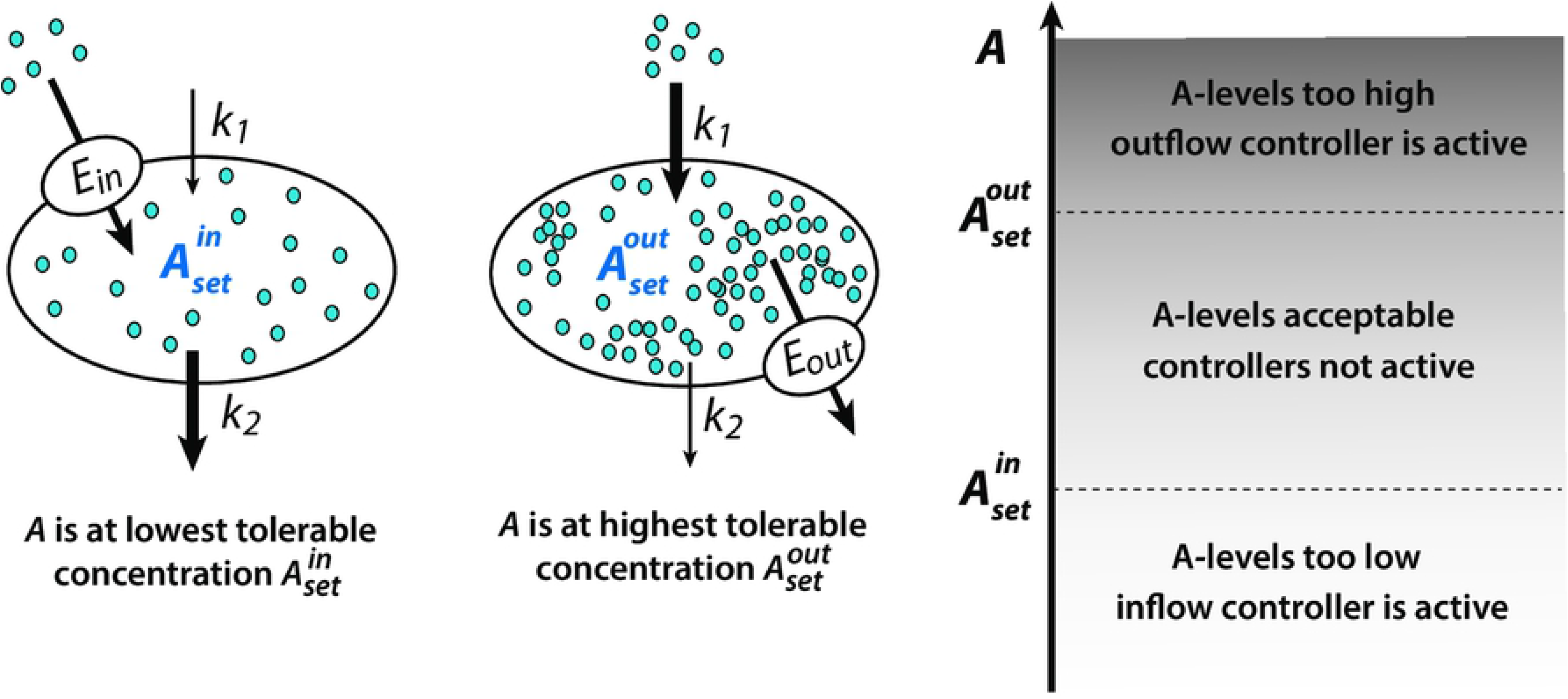
Combination of inflow/outflow controllers (indicated by the transporters *E_in_* and *E_out_*) which keep a controlled variable *A* within the controllers’ set-points, independent of the perturbation parameters *k*_1_ and *k*_2_.

Fig. 2 shows the matching controller motifs 3 (m3) and 5 (m5) [18]. Feedback structure m3 is an inflow controller, while scheme m5 is an outflow controller where *A* is the controlled variable and *k*_1_ and *k*_2_ represent perturbations.

**Fig 2.**
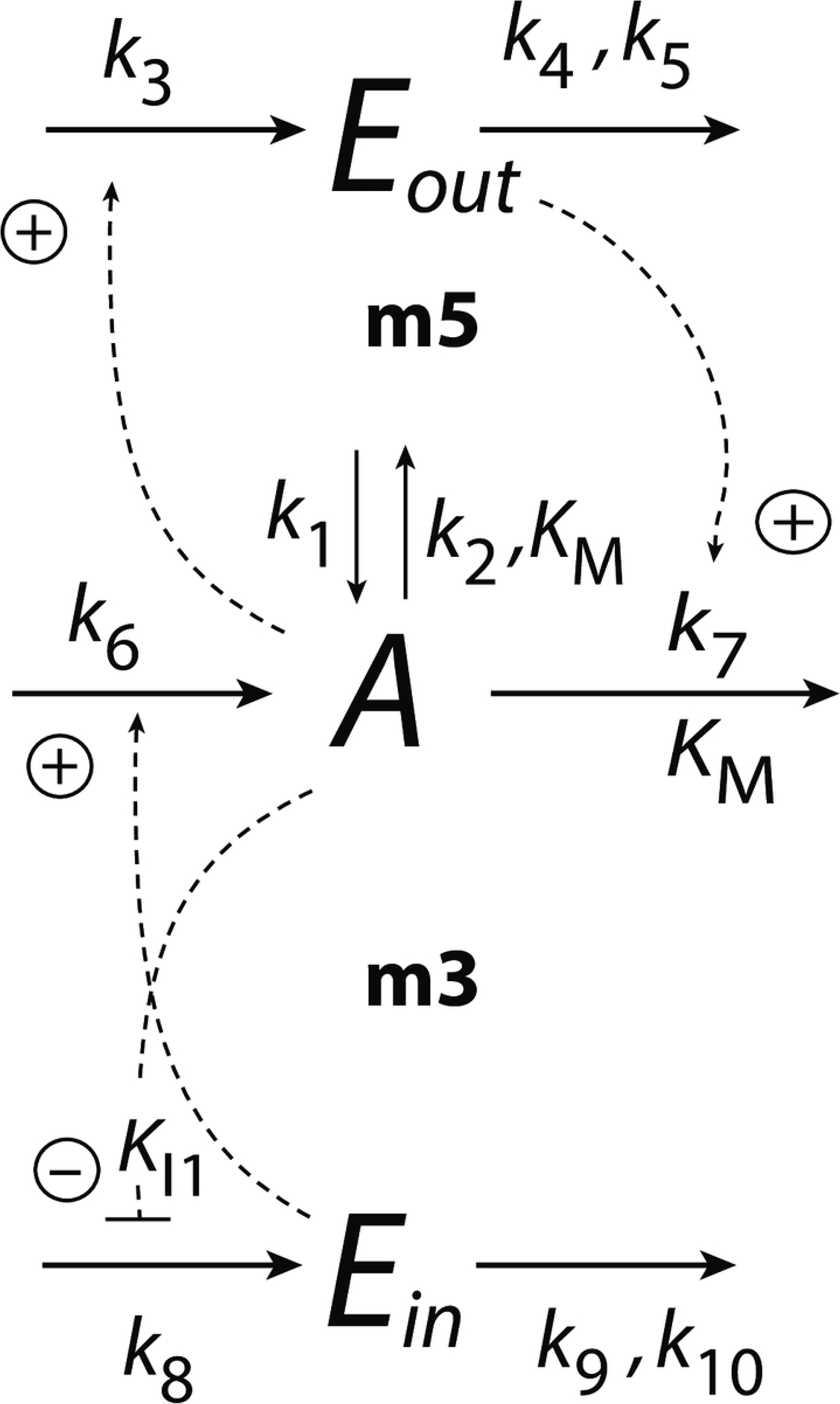
Combination of controller motifs m3 and m5 with *A* as the controlled variable. Rate parameters *k*_1_ and *k*_2_ are perturbations.

If the degradation reactions of *A* are purely first-order with respect to *A* (*K_M_* not used), the steady state concentrations of the system are non-oscillatory. In this case the rate equations for the combined controllers are:

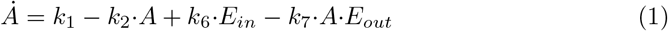

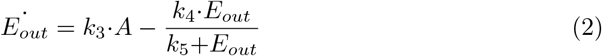

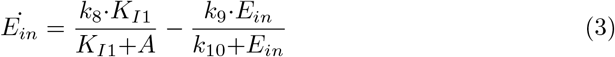

The controllers’ set-points, 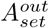 and 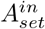, are calculated by setting 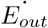 and 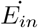 to zero. The degradation reactions with respect to *E_out_* and *E_in_* are zero-order, i.e., *k*_5_≪*E_out_* and *k*_10_≪*E_in_*, which is the requirement of getting integral feedback [18]. The set-points of the outflow and inflow controllers are then:

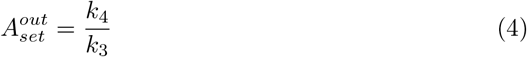

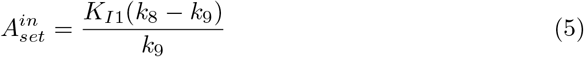

In Fig 3a the red color indicates the *A* values that are close or at the set-point of the outflow controller m5 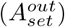, while the purple color shows the *A* values close or at 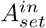. Note the corresponding up- and downregulation of *E_out_* and *E_in_* (Fig 3b).

**Fig 3.**
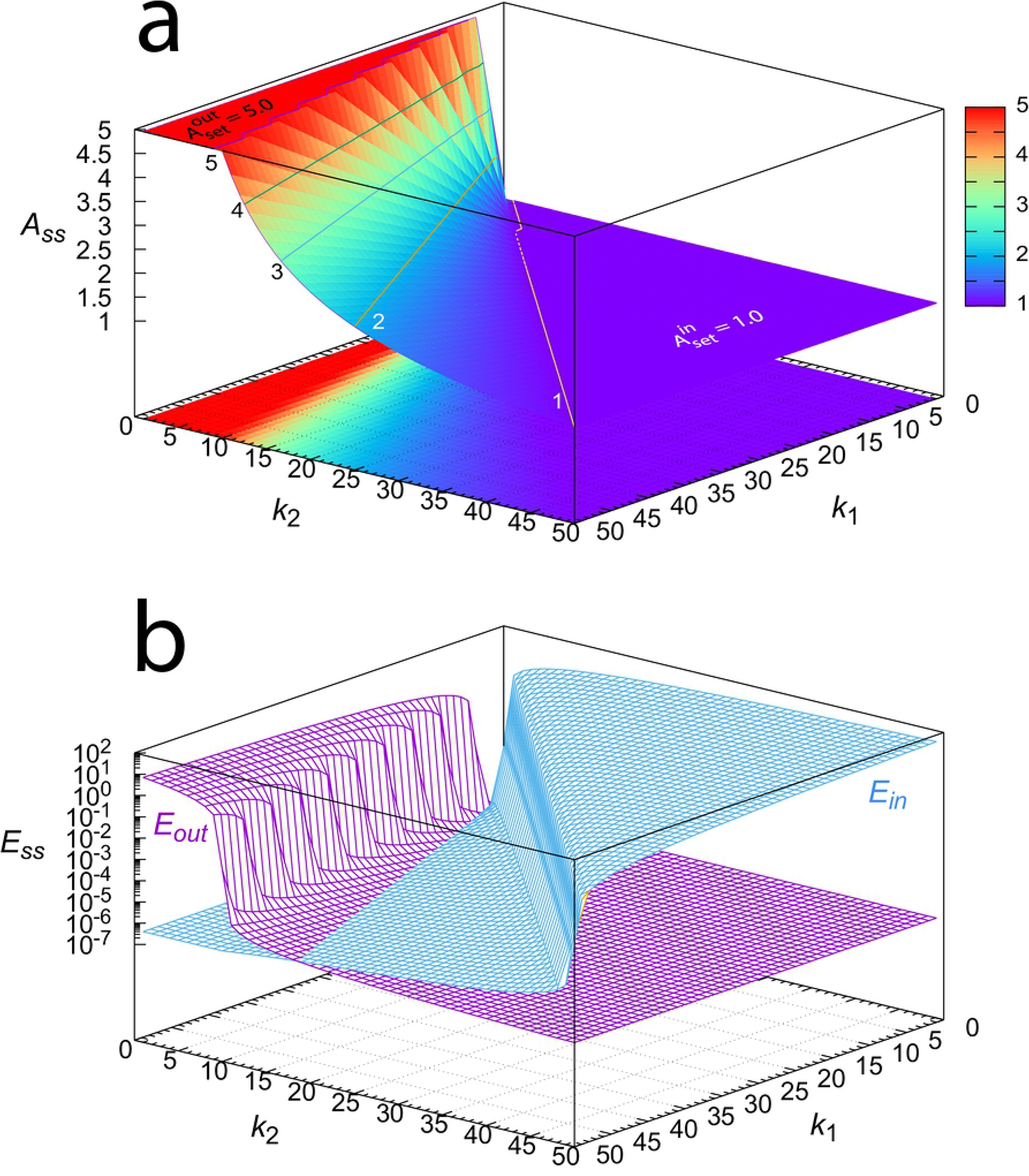
Steady state values of *A*, *E_out_*, and *E_in_* of the combined m3-m5 controllers (Fig 2) as a function of perturbation parameters *k*_1_ and *k*_2_. *k*_1_ and *k*_2_ vary between 1.0 and 50.0 with increments of 1.0. (a) Steady state values of *A*. Numbers 1-5 in the plot indicate the contour lines having this value of *A_ss_*. (b) Steady state values of *E_out_* and *E_in_*. Rate constants: *k*_3_=10.0, *k*_4_=50.0, *k*_5_=1 × 10^−4^, *k*_6_=*k*_7_=1.0, *k*_8_=2.0, *k*_9_=1.0, *k*_10_=1 × 10^−6^, *K_I_*=1.0. Initial concentrations when calculating *A_ss_* for each *k*_1_, *k*_2_ pair: *A*_0_=3.0, *E_in_*=*E_out_*=0.0; integration time: 1000 time units. For an interactive visualization of the surfaces, see S1 Gnuplot.

Fig 3 shows the steady state values of *A*, *E_out_*, and *E_in_* as a function of the perturbation parameters *k*_1_ and *k*_2_. It shows that the combined controllers in Fig 2 can keep variable *A* between the set-points of the m3 and m5 controllers.

### Oscillatory control mode

When the degradation reactions become zero-order with respect to *A* the rate equation of *A* becomes

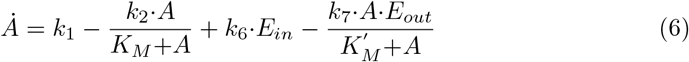

with *K_M_*, 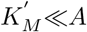. In this case both the m3 and the m5 controllers become oscillatory. For the m5 feedback loop the period can be calculated to [16]:

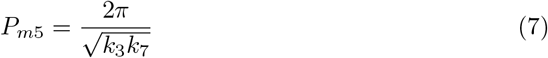

For the m3 feedback we can apply an “harmonic oscillator approximation” [28], which leads to

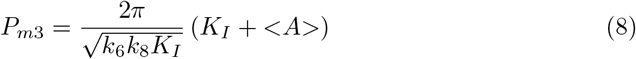

where <*A*> is the average value of *A* defined as

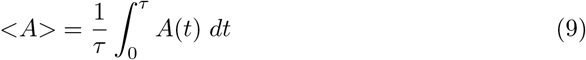

Fig 4 shows a comparison between the two combined controllers when using for *A* Eq 1 (Fig 4a) and when using Eq 6 (Fig 4b). In phase 1 the outflow perturbation *k*_2_ is largest (*k*_1_=1.0, *k*_2_=10.0) while in phase 2 the inflow perturbation is largest (*k*_1_=10.0, *k*_2_=0.0). Rate constant values have been chosen such that during phase 1 the dominating inflow controller (m3) has a period of 24 time units (Fig 4c), while during phase 2 the dominating outflow controller (m5) has a period of approximately 5 time units (Fig 4d). These rate constant values also take part in defining the set-point for the inflow controller m3 to 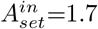 (Eq 5) and the set-point of the outflow controller m5 to 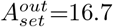 (Eq 4).

**Fig 4.**
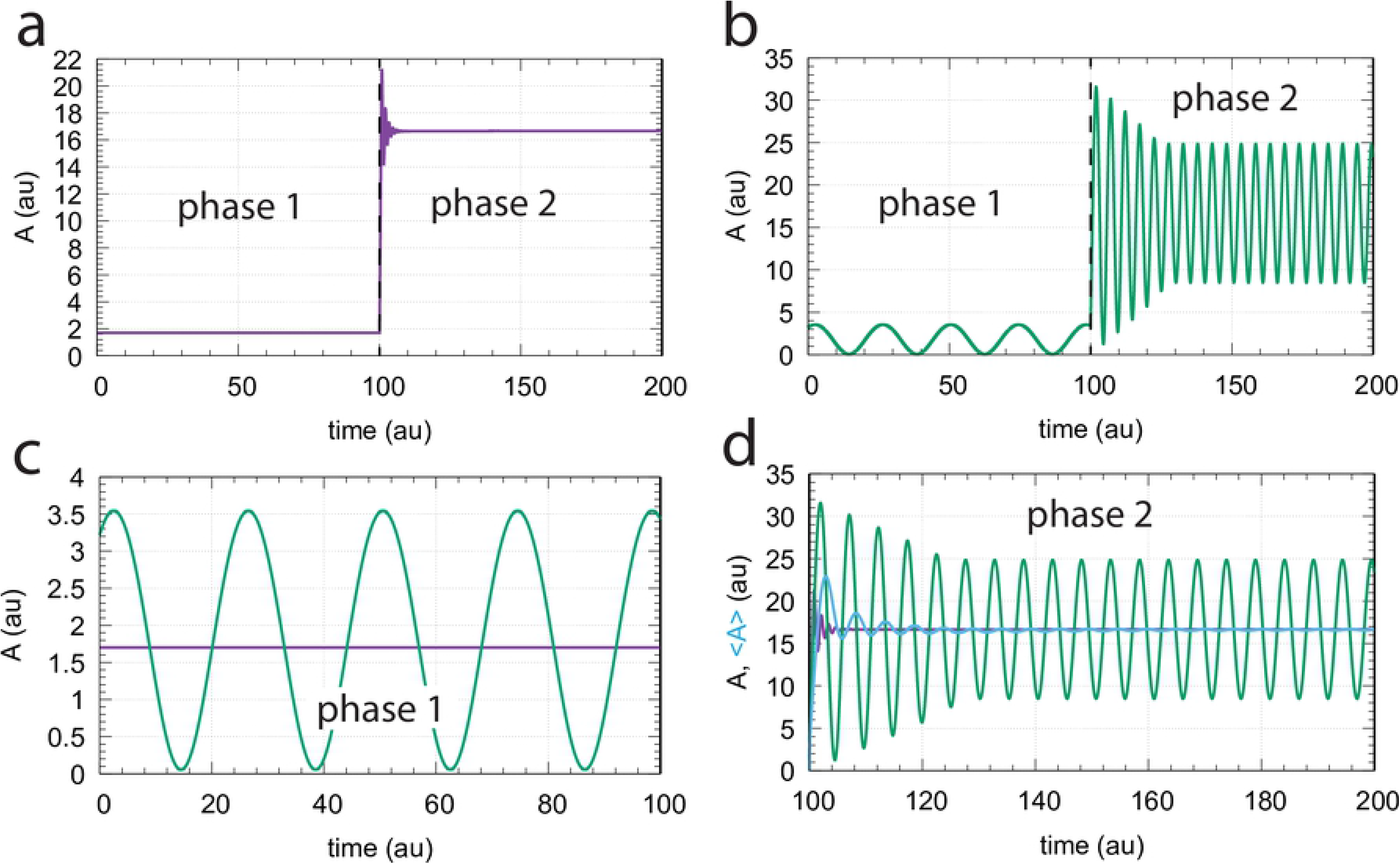
Comparison between oscillatory and nonoscillatory controller modes of combined motifs m3 and m5. (a) Nonoscillatory behavior when rate of *A* is given by Eq 1. (b) Oscillatory behavior when describing 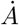 by Eq 6. (c) Comparison between oscillatory and nonoscillatory behavior in phase 1 when *k*_1_=1.0 and *k*_2_=10.0. Average value of oscillatory *A*, <*A*>, is precisely at 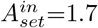. (d) Comparison between oscillatory and nonoscillatory behavior in phase 2 when *k*_1_ and *k*_2_ have changed to respectively 10.0 and 0.0. <*A*> (blue line), approaches rapidly the set-point of the outflow controller 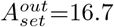. Other rate constants: *k*_3_=3.0, *k*_4_=50.0, *k*_5_=1 × 10^−8^, *k*_6_=0.7, *k*_7_=0.5, *k*_8_=1.2, *k*_9_=1.0, *k*_10_=1 × 10^−6^, *K_M_*, 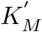 (when applied) both 1 × 10^−6^, and *K_I_*=8.5. Initial concentrations for phase 1, panel a: *A*_0_=1.700, *E*_*out*,0_=1.136 × 10^−9^, *E*_*in*,0_=2.286 × 10^1^; Initial concentrations for phase 1, panel b: *A*_0_=3.219, *E*_*out*,0_=2.395 × 10^−9^, *E*_*in*,0_=1.322 × 10^1^.

In the oscillatory control mode (Eq 6) the period of the dominant (ruling) controller is established (Fig 5a). The amplitude of the inflow controller m3 is practically constant while for outflow controller m5 the amplitude increases. However, the amplitude of the *A*-oscillations ceases completely when *k*_1_ and *k*_2_ are equal (Fig 5b). Fig 5c shows that the oscillatory controllers follow (on average) closely the controllers set-points of the nonoscillatory state (Eqs 4 and 5). Fig 5d shows the increase of the average values of respectively *E_in_* and *E_out_* (<*E_in_*> or <*E_out_*>) when *k*_2_ or *k*_1_ become the dominating perturbations, respectively.

**Fig 5.**
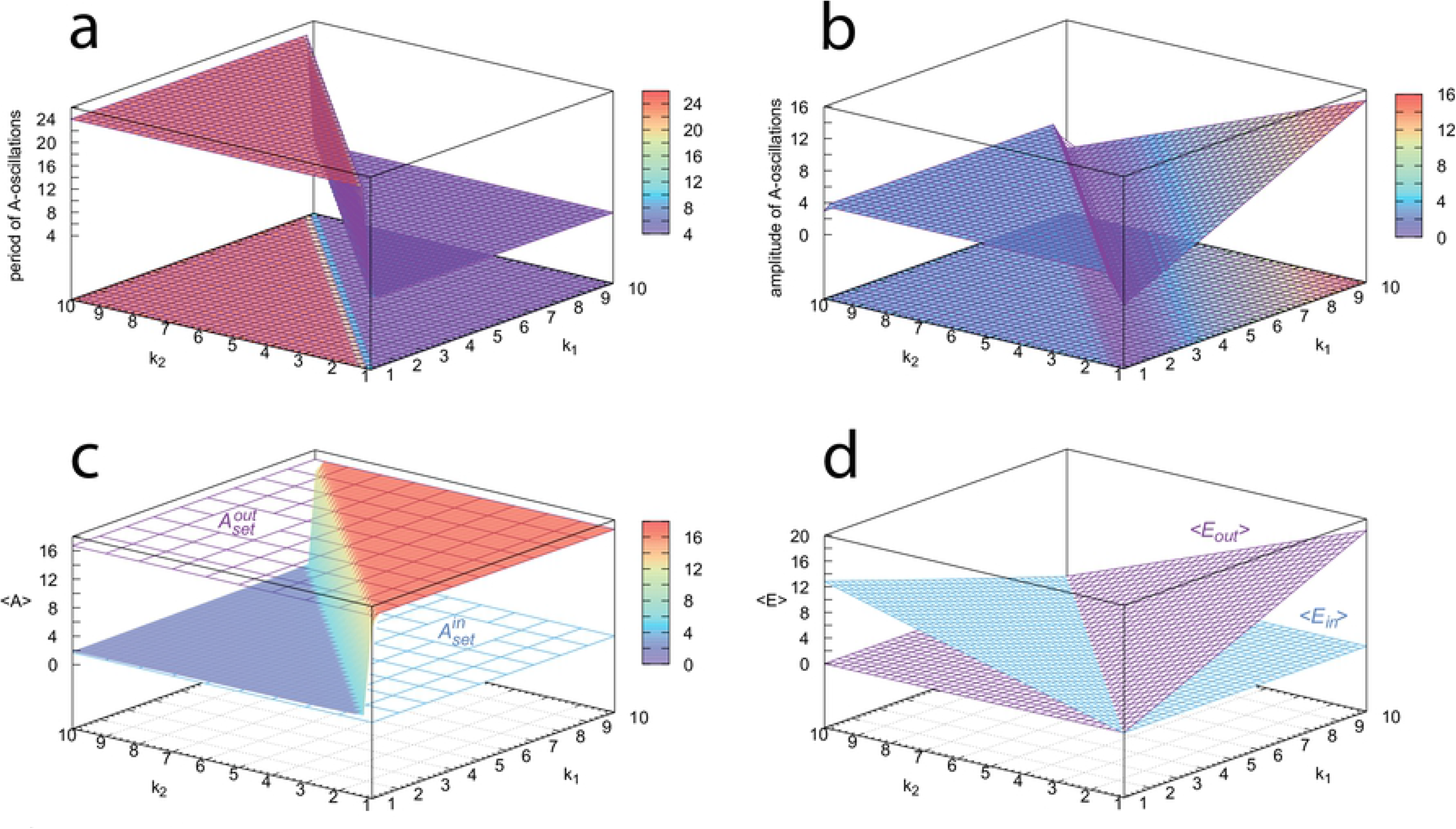
Overview of the oscillatory properties of the controllers (Fig 4). (a) The period switches in dependence whether the inflow or outflow controller is dominating. (b) Amplitude of the *A*-oscillations. (c) The controllers’ <*A*> values follow closely their set-points. (d) Average values of the oscillatory *E_in_* and *E_out_* concentrations and their up-regulation in dependence of the pertrubations. Rate constants and initial concentrations as for the oscillatory case in Fig 4. All properties were calculated after 500 time units when (oscillatory) steady state conditions were established. See also S2 Gnuplot for an interactive visualization for each panel.

### p53 regulation by inflow and outflow control

p53 is a protein often described as the “guardian of the genome” [34]. p53 takes part in cell fate decisions [35] with respect to internal or environmental disturbances and is involved in cell cycle arrest. p53 is also considered to prevent tumor development by inducing apoptosis in response to DNA-damage and other stress signals [36].

In normal unstressed cells p53 is at low levels due to different proteasomal degradation reactions including ubiquitin-dependent [37] and ubiquitin-independent [38–40] pathways. For the ubiquitin-independent pathways the enzyme NAD(P)H:quinone oxidoreductase 1 (NQO1) has been indicated to have a major regulatory role [41,42]. In unstressed cells there is further evidence that p53 and the circadian clock [30] undergo cooperative regulations [43–46] via the Per2 protein. Per2 is not only an important component of the human circadian oscillator [47], but also takes part in the input and output pathways of the clock [48]. Under normal (unstressed) conditions p53 has been found to inhibit expression of Per2 by binding to its promotor [43]. Overexpressing Per2 in HCT116 cells resulted in a significant increase in p53 mRNA [43] or in an induced apoptosis in lung cancer cells [49], indicating that Per2 can activate the synthesis of p53. In addition, Per2 has been found, by its binding to p53, to inhibit the Mdm2-mediated degradation of p53 and thereby stabilizing p53. This Per2-p53 feedback loop has the typical properties of an inflow type of controller and is of the same basic structure (motif 3 [18]) as the *A*-*E_in_* loop in Fig 2. This suggests that p53 is kept by the circadian clock at an acceptable minimum (“preconditioning” [45]) level with set-point p53_min_, which allows a sufficiently rapid up-regulation of p53 in the case of stress/DNA-damage.

In case of stress/DNA-damage p53 is up-regulated by ataxia telangiectasia mutated (ATM) kinase [50,51]. The treatment of human MCF7/U280 cell lines with 10 Gy gamma radiation showed oscillations in p53 and Mdm2 with a period length of about 5-6 hours [52]. An interesting feature of these oscillations is that their period is relatively stable, while there is a considerable variation in their amplitude. It has also been pointed out [52] that a significant fraction of the MCF7/U280 cells (about 40% at 10 Gy) do not oscillate, i.e., either showed no variations in p53 or showed only slowly varying fluctuations. In analyzing the p53-Mdm2 negative feedback loop, Jolma et al. [53] found that the loop can show harmonic oscillations when the respective degradations of p53 and Mdm2 approach zero-order. The conservative feature of these oscillations not only could explain the constancy of the period and the stochastic variation in the amplitude, but as a motif 5 outflow controller [18], the set-point of the p53-Mdm2 loop provides an upper p53 concentration limit, probably to avoid a premature apoptosis of cells.

A Fourier analysis of the p53 oscillations [54] showed indeed a major harmonic peak at about 5-6h along with minor 2nd and 3rd-order harmonics at lower periods. The rise of the Fourier transform at higher period lengths (>10h) provides evidence for an additional loop, which Geva-Zatorsky et al. [54] considered to be a feedback loop between ATM and p53. In this feedback loop the active (phosphorylated) form of ATM (ATM*) activates p53 via CHK2 (checkpoint kinase 2) [51,55], while p53 inhibits ATM* via the activation of phosphatase WIP1 [51,56,57]. A closer look at the p53-ATM* loop shows that it acts as a motif 1 ([18]) inflow controller. The inflow control function of this loop suggests that the loop’s set-point, 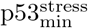, keeps the p53 concentration in stressed cell at a minimum level, but higher than the set-point imposed by the circadian clock. As we will show below the set-point defined by the ATM* controller increases with the stress level activating ATM and counteracts perturbations which may accidentally drive p53 to lower levels.

Based on these observations we arrive at a p53 homeostatic model of three interlocked feedback loops with period lengths of p53 oscillations which are dependent on the stress level and the actual feedback loop responding to it. Fig 6 shows a schematic representation of the model. In unstresssed cells the inflow control properties of the p53-Per2 feedback loop, analogous to motif 3, ensures that p53 is on average at a minimum low level (with set-point p53_min_) compensating for the proteasomal ubiquitin-independent degradations of p53 via NQO1. In stressed cells several factors increase the level of p53, including its activation by ATM, the inhibition of the ubiquitin-independent degradation pathways of p53, and the stabilization of p53 by chaperones such as HSP90. The inflow control structure between p53 and the activated ATM loop (motif 1) now drives p53 levels up to 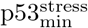. As stabilization and concentration of p53 further increases, the Mdm2-p53 control loop (motif 5) will oppose further increase of p53, at least temporarily. However, since the set-point of this controller is given by the ratio between synthesis and degradation rates of Mdm2, the set-point of the Mdm2-controller may increase when Mdm2 is stabilized by chaperones/HSP90 and the Mdm2 turnover is inhibited [58].

**Fig 6.**
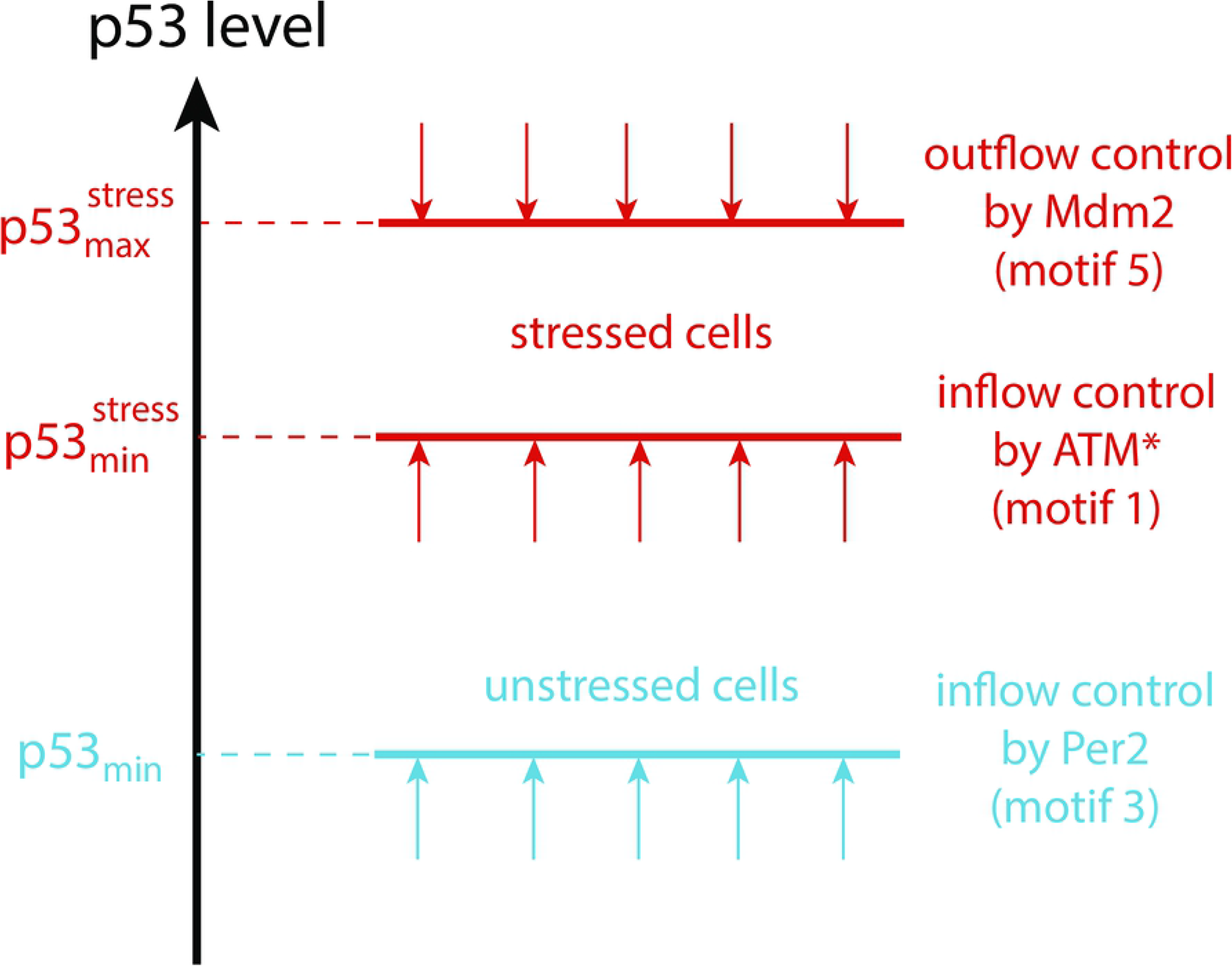
Suggested inflow/outflow regulation of p53 for unstressed and stressed cells. In unstressed cells the p53-Per2 loop keeps p53 at the minimum level p53_min_ (indicated by the blue arrows), opposing the dominant degradation reactions of p53 by Mdm2-independent proteasomal degradation. When stress is recognized, the active ATM form, ATM*, is up-regulated and the ATM*-p53 loop drives p53 concentrations to 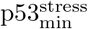 (indicated by red upright arrows). Indicated by the red downward pointing arrows, the p53-Mdm2 feedback loop with set-point 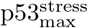 assures that cells are not prematurely driven into apoptosis.

Fig 7 shows the reaction scheme of the model for unstressed and stressed cells. The model consists of 9 coupled rate equations. Dependent on the stress level, reactions outlined in light gray are low in their reaction rates/concentrations, while reactions outlined in black are the dominating ones. For the sake of simplicity we used the single variable *k*_30_ to mediate the stress into the network, both for the activation of ATM (with activation constant *K_as_*) and the inhibition of the NQO1-mediated proteasomal degradation of p53 (with inhibition constant *K_Is_*). In addition, we also include a stress-related increase of p53 via *k*_1_ in parallel to its activation by ATM*. This additional activation may be related due to the presence of reactive oxygen species [59], although the reaction pathways, as indicated by the question mark, appear to be not well understood.

**Fig 7.**
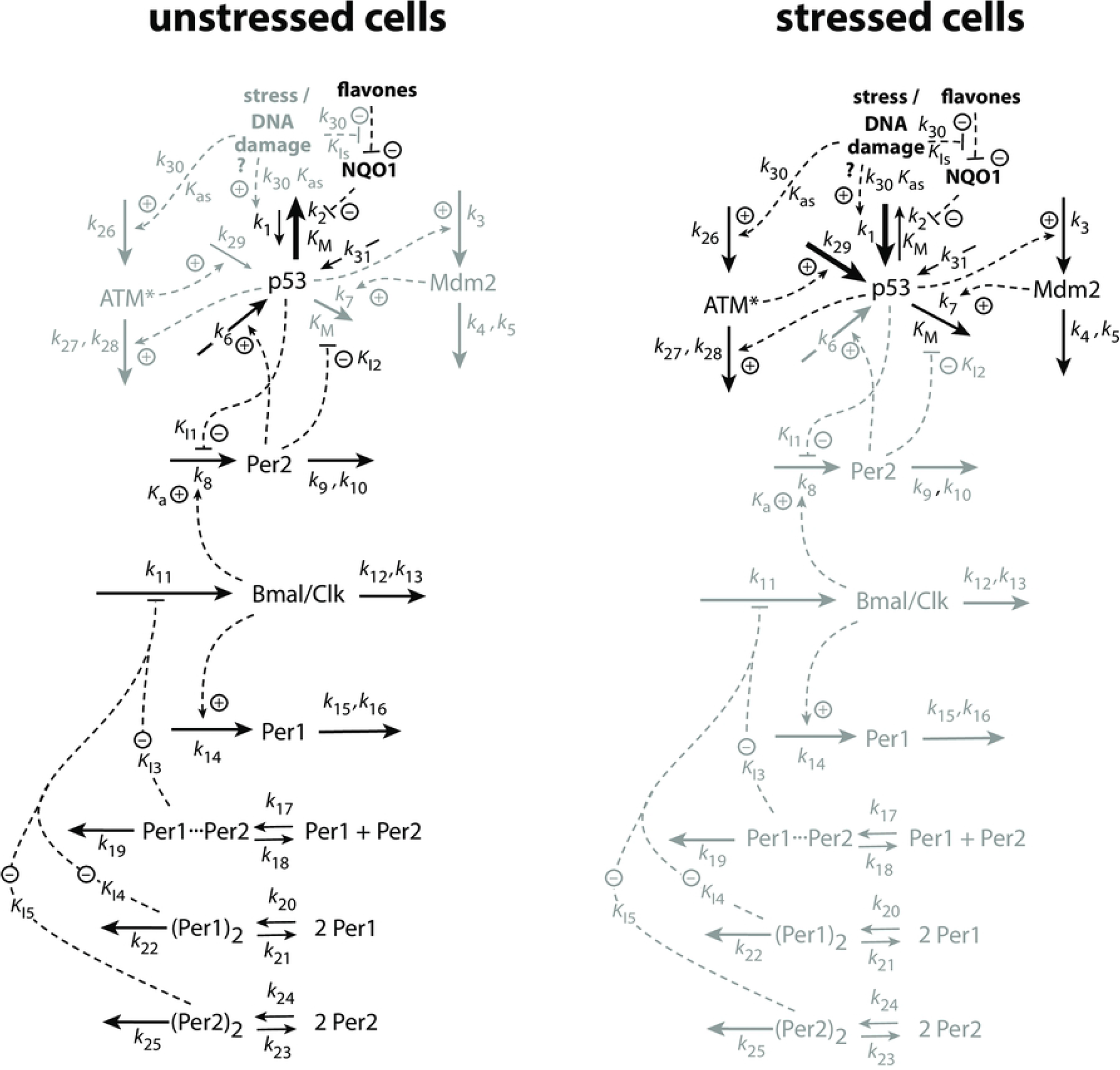
Model of the p53 regulatory feedback loops. Dependent on the stress level *k*_30_, certain parts of the network become down-regulated (outlined in gray), while other parts are up-regulated (indicated in black). For rate equations, see main text.

The activation of ATM to ATM* by the stress level *k*_30_ is described by the rate equation

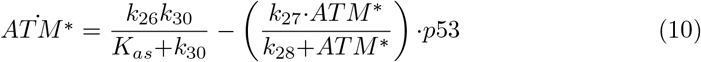

The rate equation for *p*53 consists of four inflows and two outflows (Fig 7).

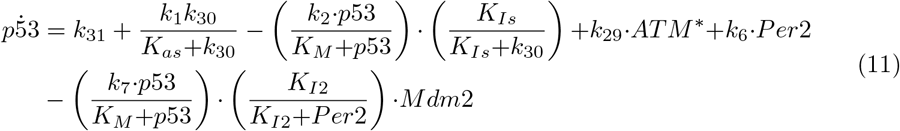

The first term, *k*_31_, is a constitutive (constant) expression term for p53 [60], while the second and third terms represent, respectively, stress-induced increase of p53 production and a stress-induced inhibition of the proteasomal ubiquitin-independent degradation of p53 via NQO1.

The other rate equations are:

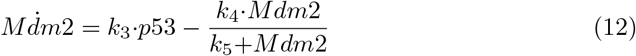

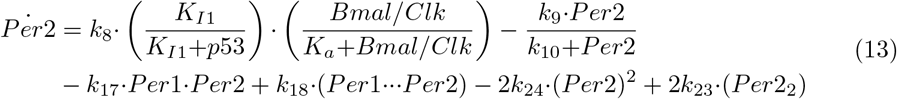

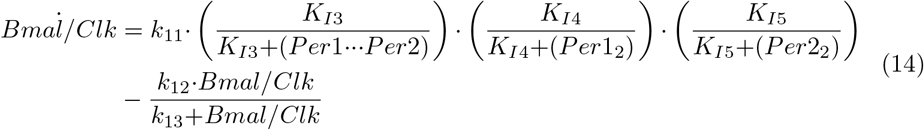

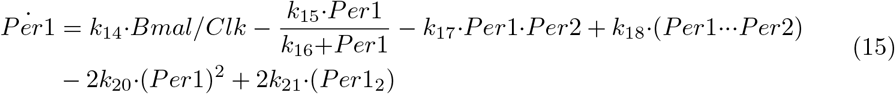

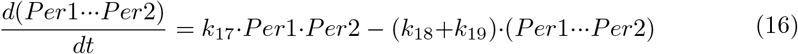

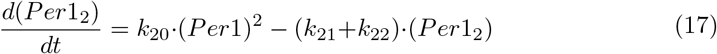

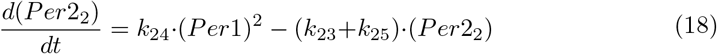

In the above equations (*Per*1_2_) and (*Per*2_2_) denote the respective concentrations of the *Per*1 and *Per*2 homodimers, while (*Per*1⋯*Per*2) denotes the concentration of the *Per*1-*Per*2 heterodimer.

As indicated by Fig 6 p53 is controlled in this model by three feedback loops. Rate parameters have been chosen such that each of the feedback loops has integral control/feedback with defined set-points and, when oscillatory, defined period lengths.

In the absence of stress p53 is rapidly degraded by the proteasome. In this case Per2 acts as an inflow controller with a set-point given by Eq 5, i.e.,

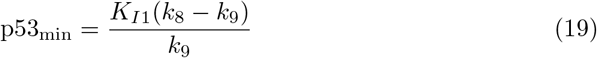

Since we assume that the degradation reaction of p53 are zero-order with respect to p53, the Per2 controller oscillates around p53_min_ with a period described by Eq 8, i.e.,

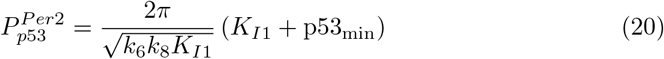

The values of *k*_6_ (0.7), *k*_8_ (1.2), *K*_*I*1_ (8.0), and *k*_9_ (1.0) have been chosen such that p53_min_ is relatively low, i.e., 1.6. 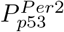 is thereby in the circadian range (≈24h).

When stress is present, but not too high (0.1 ≤ *k*_30_ ≤ 1), ATM* determines the average concentration of p53 and the frequency of the p53 oscillations. By setting Eq 10 to zero and assuming zero-order degradation of ATM* with respect to ATM* the set-point of this controller is dependent on the stress level *k*_30_, i.e.,

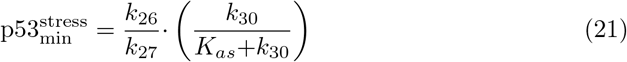

Rate parameters *k*_26_ (90.0), *k*_27_ (10.0), and *K_as_* (3.0) have been chosen such that 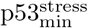 has a maximum value of 9.0 when *k*_30_ is high. This value has been arbitrarily chosen, with the only requirement that 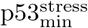 should be higher than p53_min_. The ATM* controller’s period (being a motif 1 controller) is calculated to be (S1 Text):

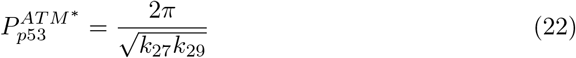

Using, rather arbitrarily *k*_29_=1.0, the period of the ATM* controller is approximately 2h.

For high stress levels (*k*_30_ > 1) the Mdm2 outflow controller keeps p53 at a much higher set-point analogous to Eq 4, i.e.,

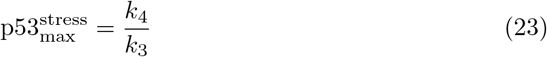

Ignoring the influence of noise [16], p53 oscillates now around the set-point described by Eq 23 with period

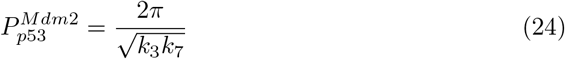

Using values of *k*_3_ and *k*_7_ of respectively 3.0 and 0.5 the period of the Mdm2 controller is 5.1h. With *k*_4_=50.0 the value of 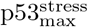 is 16.6 and defines an upper bound of the p53 concentration. How this upper bound can be further increased and finally leads to apoptosis will be discussed below.

Fig 8 shows how the steady state period and average levels of p53 change with the stress signal *k*_30_. At certain stress levels the controllers Per2, ATM*, and Mdm2 are individually up-regulated. They defend their set-points and frequencies of the p53 oscillations. Mrosovsky [61] termed the defense of different environmentally-induced set-points as “reactive rheostasis”.

**Fig 8.**
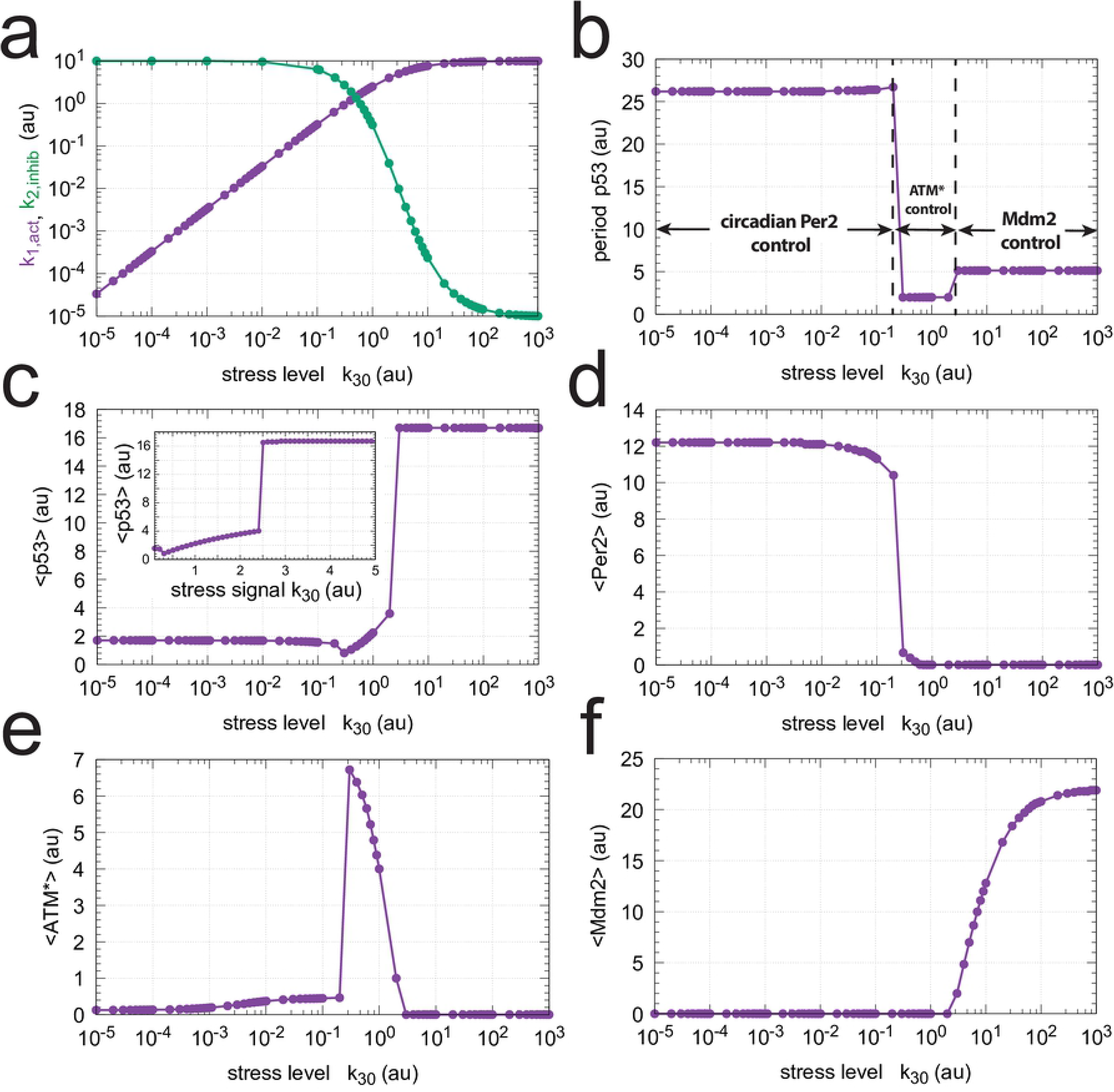
Change of steady state levels in the model (Fig 7) as a function of stress parameter *k*_30_. (a) Change of *k*_1,*act*_ (Eq 25) and *k*_2,*inhib*_ (Eq 26), (b) p53 period length, (c) average p53 concentration <p53>; inset: same figure, but abscissa (*k*_30_) is linear. (d) average Per2 concentration <Per2>, (e) average ATM* concentration <ATM*>, and (f) average Mdm2 concentration <Mdm2>. Parameter values: *k*_1_=*k*_2_=10.0, *k*_3_=3.0, *k*_4_=50.0, *k*_5_=1 × 10^−6^, *k*_6_=0.7, *k*_7_=0.5, *k*_8_=1.2, *k*_9_=1.0, *k*_10_=1 × 10^−6^, *k*_11_=0.85, *k*_12_=0.7, *k*_13_=1 × 10^−6^, *k*_14_=1.0, *k*_15_=0.7, *k*_16_=1 × 10^−6^, *k*_17_=*k*_18_=1 × 10^3^, *k*_19_=0.0, *k*_20_=10.0, *k*_21_=0.5, *k*_22_=0.0, *k*_23_=1 × 10^6^, *k*_24_=1 × 10^3^, *k*_25_=0.0, *k*_26_=90.0, *k*_27_=10.0, *k*_28_=1 × 10^−6^, *k*_29_=1.0, *k*_31_=1.0, *K_M_*=1 × 10^−6^, *K*_*I*1_=*K*_*I*3_=*K*_*I*4_=*K*_*I*5_ =8.0, *K*_*I*2_=1 × 10^6^, *K_Is_*=3.0, *K_a_*=0.012, *K_as_*=3.0. Initial concentrations: p53_0_=1.49, Mdm2_0_=9.84 × 10^−8^, Per2_0_=3.82 × 10^−2^, Bmal/Clk_0_=1.00, Per1_0_=1.48 × 10^−1^, Per1⋯Per2_0_=5.66 × 10^−3^, (Per1)_2_, 0=1.82 × 10^−1^, (Per2)_2_, 0=1.46 × 10^−6^, 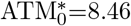. Simulation time 3000 time units (h).

In Fig 8a we suggest how a stress-induced inflow to p53 (Eq 25) and a stress-induced inhibition of p53-degradation (Eq 26) may change with stress level *k*_30_.

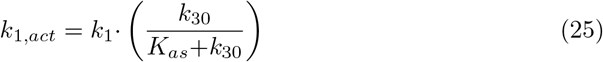

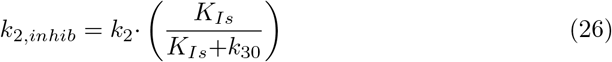

Panels b and c show how the individual controllers, dependent on the stress level, determine p53’s period length and average level. Panels d-f show the controllers Per2 (d), ATM* (e), and Mdm2 (f) and their appearances at different stress levels.

Fig 9 shows the steady state oscillations at four different stress levels. Panel a shows the Per2 and p53 oscillations at low/no stress. In agreement with experiments [45] Per2 peaks a couple of hours earlier than p53. As indicated in Fig 8a Per2 has control over p53 rhythmicity and level in unstressed cells. In panel b, at minor stress levels, we see the Per2 and p53 oscillations at the transition to ATM* control, which is indicated by the appearance of the short period oscillations of the ATM* controller. Panel c shows the oscillations when the ATM* concentration is relatively high (Figs 8e), while in panel d the Mdm2 controller has taken over and is determining the level and period of the p53 oscillations. (Figs 8f).

**Fig 9.**
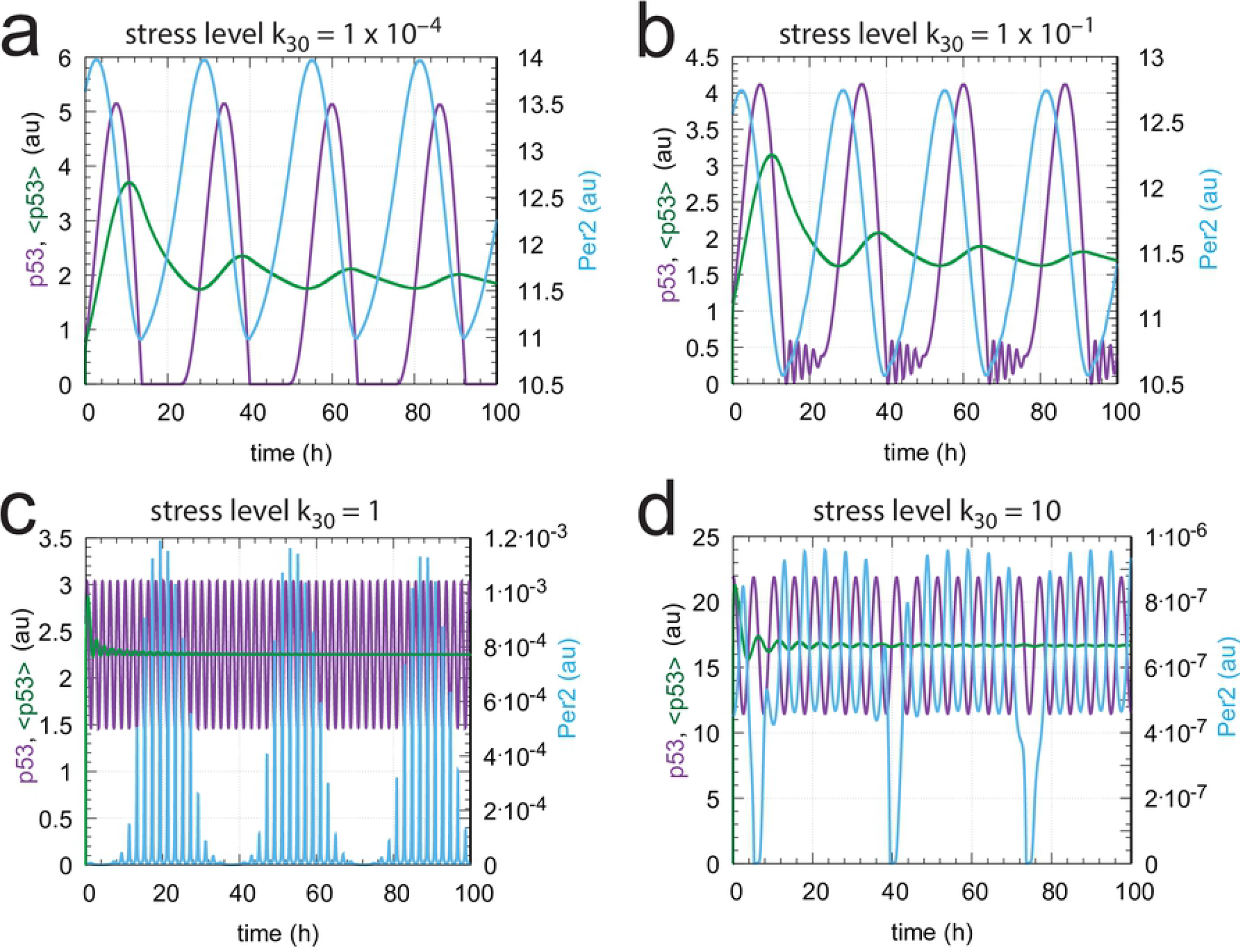
Steady state oscillations of p53 and Per2, together with <p53>, at different stress levels *k*_30_. Parameter values are the same as in Fig 8. (a) Oscillatory behavior when *k*_30_=1 × 10^−4^. Per2 determines the state of p53. Initial concentrations: p53_0_=7.65 × 10^−1^, Mdm2_0_ =4.82 × 10^−8^, Per2_0_=1.36 × 10^1^, Bmal/Clk_0_=0.39, Per1_0_=7.34 × 10^−2^, (Per1⋯Per2)_0_=1.00, (Per1_2_)_0_=2.21 × 10^−1^, (Per2_2_)_0_=1.86 × 10^−1^, 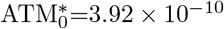. (b) *k*_30_=1 × 10^−1^. The high frequency oscillations of the ATM* controller begin to appear, but the Per2 controller still determines p53 period length. Initial concentrations: p53_0_=1.11, Mdm2_0_=7.12 × 10^−8^, Per2_0_=1.26 × 10^1^, Bmal/Clk_0_=0.47, Per1_0_=8.44 × 10^−2^, (Per1⋯Per2)_0_=1.06, (Per1_2_)_0_=2.35 × 10^−1^, (Per2_2_)_0_=1.58 × 10^−1^, 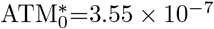. (c) *k*_30_=1.0. The ATM* controller has taken over and p53 oscillates with a period of 2h (Eq 22) around the controller’s set-point 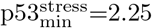 (Eq 21). Initial concentrations: p53_0_=2.58, Mdm2_0_=1.83 × 10^−7^, Per2_0_=5.56 × 10^−6^, Bmal/Clk_0_=0.17, Per1_0_=3.55 × 10^−1^, (Per1⋯Per2)_0_=1.98 × 10^−6^, (Per1_2_)_0_=3.03, (Per2_2_)_0_=3.10 × 10^−14^, 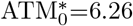. (d) *k*_30_=10.0. Mdm2 is the dominant controller. p53 oscillates with a period of 5.1h (Eq 24) around a set-point of 16.6 (Eq 23). Initial concentrations: p530=21.85, Mdm2_0_=1.17 × 10^1^, Per2_0_=4.54 × 10^−7^, Bmal/Clk_0_=0.41, Per1_0_=4.03 × 10^−1^, (Per1⋯Per2)_0_=1.83 × 10^−7^, (Per1_2_)_0_=3.52, (Per2_2_)_0_=2.06 × 10^−16^, 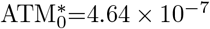.

Each of the three controllers, Per2, ATM*, and Mdm2, defend their set-points. As inflow controllers Per2 and ATM* will compensate for outflow perturbations, for example by an accidental increase of *k*_2_, while the Mdm2 controller will compensate any inflow to p53.

As an example we show the homeostatic behavior of the ATM* controller. The set-point of the ATM* controller, which depends on the stress level *k*_30_ (Eq 21) is defended towards changes in p53 outflow. Fig 10a shows the behavior of the p53 oscillations when *k*_30_=1.0 (Fig 9c) and *k*_2_ undergoes a perturbation at t=20h from 10.0 to 50.0. The set-point of the controller (2.25) is defended by an up-regulation of ATM*. Also the period (taken here arbitrarily as 1.99h, Eq 22) is kept constant (Fig 10b). Fig 10c shows the circadian oscillations of Per1 and Bmal/Clk, which are unaffected by the perturbation in *k*_2_ and keep a phase relationship in agreement with experimental results [62].

**Fig 10.**
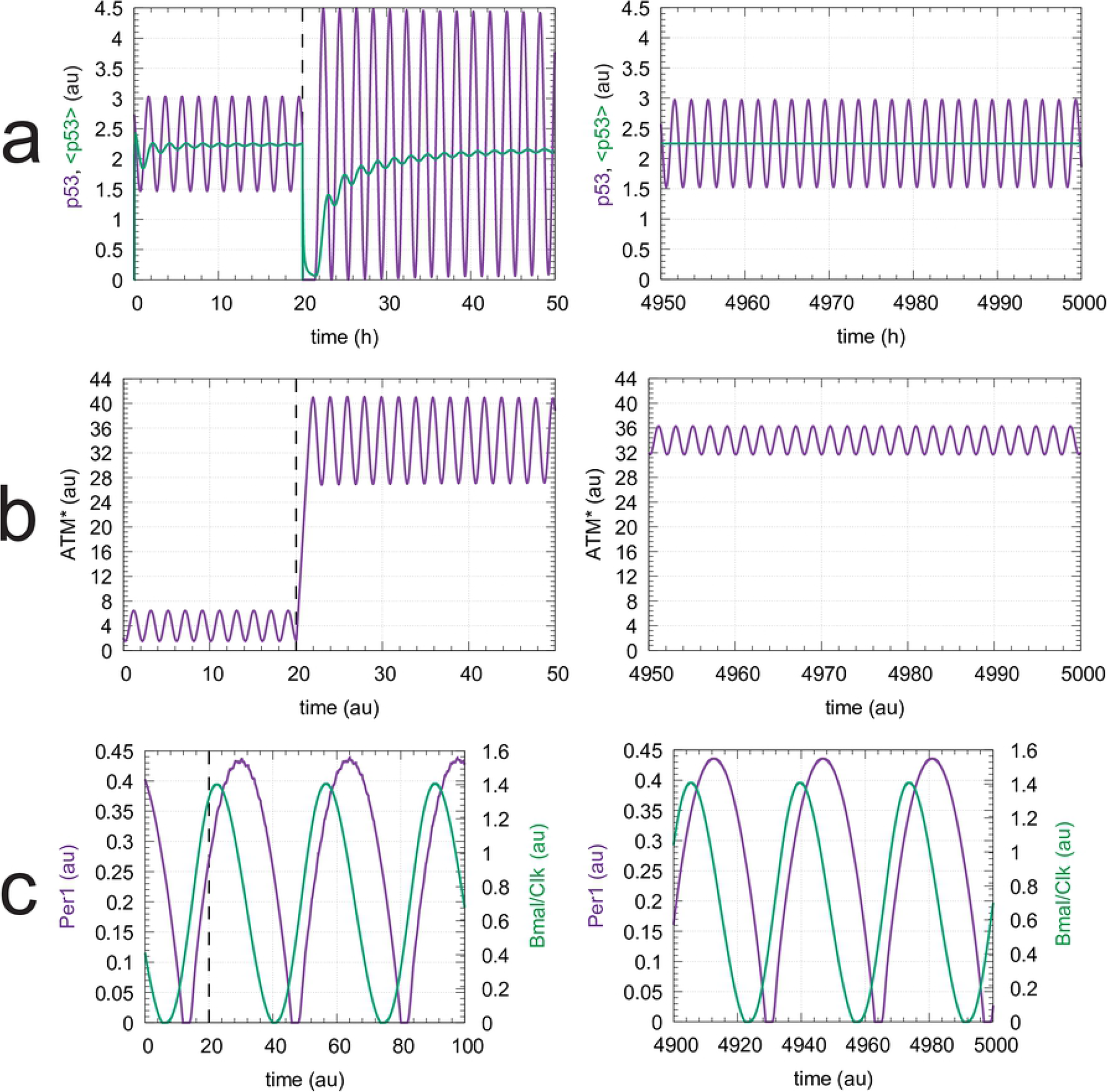
The ATM* controller defends its *k*_30_-dependent set-points (Eq 21) towards outflow perturbations. Row a, left panel: p53 oscillations and average p53 levels, <p53>, as a function of time with *k*_30_=1.0. At time t=20h *k*_2_ is increased from 10.0 to 50.0. Row a, right panel: average p53 concentration is back at the controller’s set-point (2.25) with an unchanged period length of 1.99h. Row b, left and right panels: Up-regulation of ATM* due to the change in *k*_2_. Row c, left and right panels: oscillations in Per1 and Bmal/Clk are unaffected by the *k*_2_ perturbation. Initial concentrations and rate parameters as in Fig 9c and Fig 8.

## Synergy conditions for coupled feedback loops

There are certain requirements that need to be met such that a set of coupled negative feedback motifs will cooperate and work together. As pointed out in [18] a cooperative interaction between a set of negative feedback loops (m1-m8) will depend on how the set-points of the individual controllers (determined by their 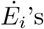) are positioned relative to each other within the concentration space of the controlled variable A. Fig 11 shows the sign structures of the 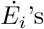 for the eight feedback structures m1-m8.

**Fig 11.**
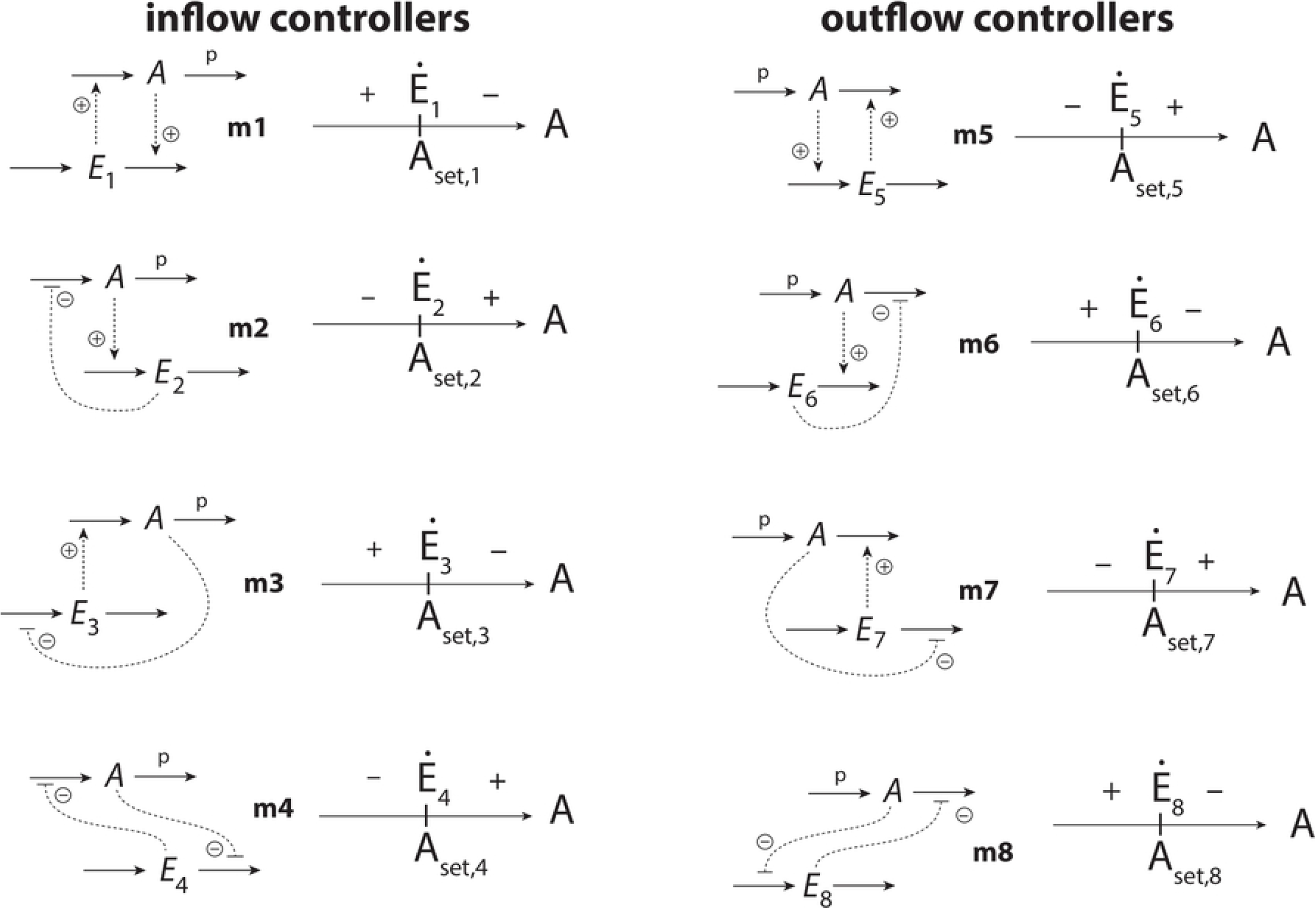
*E_i_* sign structures of the eight controller motifs m1-m8 [18]. A is the controlled variable and the *E_i_*’s are the controller species. Inflow controllers compensate outflow perturbations of *A*, while outflow controllers compensate for inflow perturbations. The perturbations acting on *A* are indicated by a “p”. For step-wise perturbations and assuming the presence of integral control, the set-point *A_set,i_* of controller *i* is determined by the steady state condition 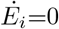.

For example, when a m1 and a m5 controller (Fig 12a) are coupled such that 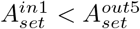 the controllers will cooperate and either m1 or m5 will dominate dependent on the perturbation acting on *A*. However, when 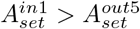 the two controllers will always work against each other, as indicated in Fig 12b. Both controllers are always in an “on-state” with the effect that *E*_1_ and *E*_5_ will increase continuously, a situation termed in control engineering as *integral wind-up* [63]. For details, see S2 Text.

**Fig 12.**
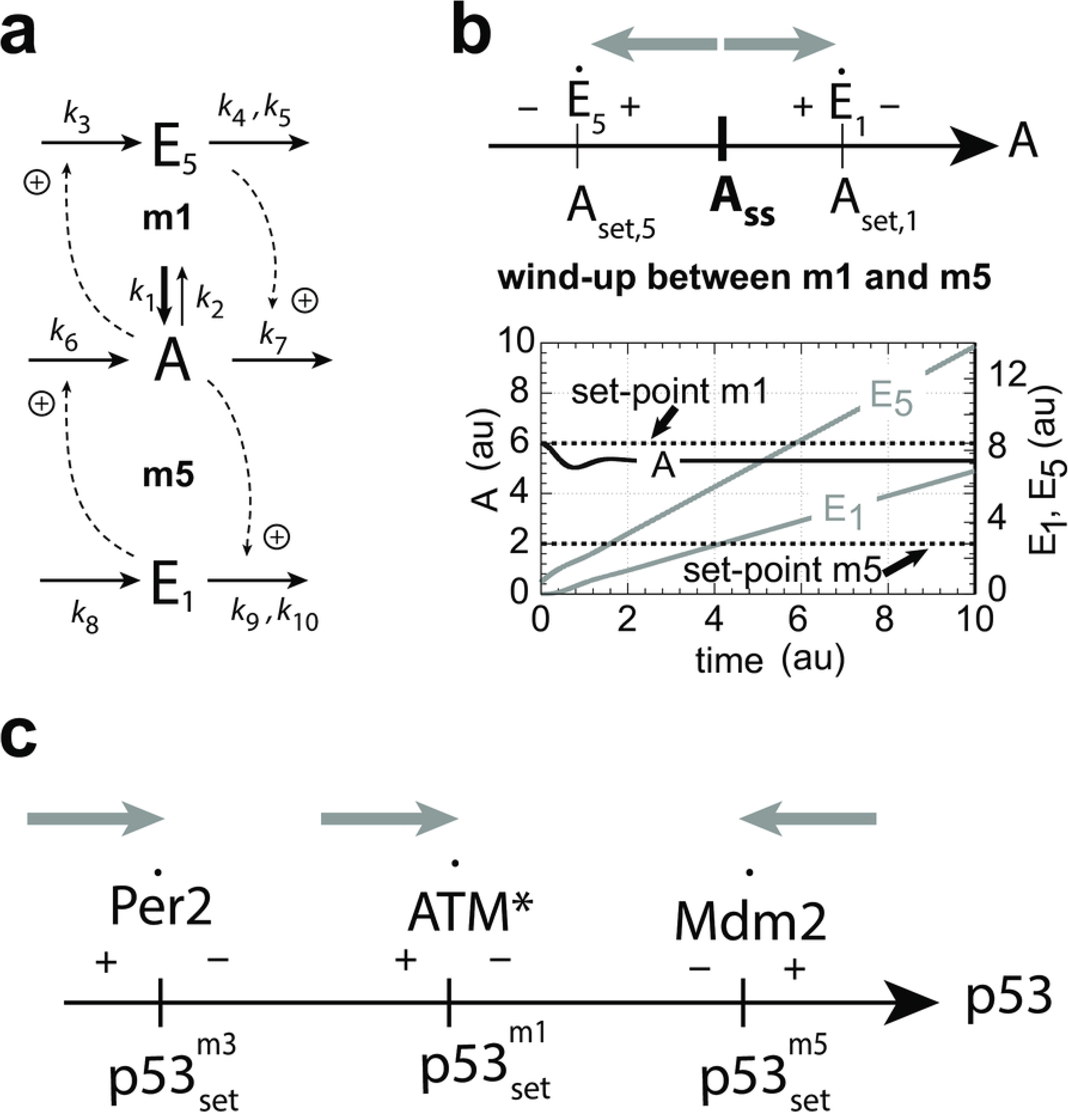
Cooperative and dysfunctional wind-up behavior in combined negative feedback loops. (a) Combined controllers m1 and m5 (S2 Text). (b) Wind-up behavior when *A*_*set*,1_ > *A*_*set*,5_. Because 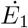 and 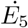 meet each other in this case with positive signs, each controller pulls in the direction of its own set-point (gray arrows). As a result, *E*_1_ and *E*_5_ increase continuously and the controlled variable *A* lies somewhere between the two set-points dependent on the individual aggressiveness [18] of m1 and m5. S2 Text shows in addition the cooperative behaviors of the two controllers when set-points are switched. (c) Cooperative behavior of the Per2, ATM*, and Mdm2 controllers with respect to p53 regulation. Gray arrows show the direction in p53-concentration space into which each controller pulls.

The structures of the interacting negative feedback loops shown in Fig 6 and Fig 7 have been taken from the literature (see references cited above). Fig 12c shows the relative setting of the set-points in the p53 concentration space for the Per2, ATM*, and Mdm2 controllers and their sign structures. It appears encouraging that these feedback loops can be placed in a naturally occurring order in p53 concentration space without wind-up.

## Roles of the individual feedback loops

The spacing of the controllers in Fig 12c suggest that the three feedback loops have certain functions in the regulation of p53. As an inflow controller, the Per2-p53 feedback loop has apparently the function to keep p53 in unstressed cells at a certain minimum level in order to allow a sufficiently rapid p53 up-regulation [44] in case DNA damage occurs.

In case of DNA damage, the ATM*-p53 loop, is up-regulated. This loop keeps p53 at a higher set-point dependent on the stress level *k*_30_. The ATM* controller defends its set-point towards increased degradations of p53 as long as stress is encountered. This suggests that as long as DNA-stress is present, the ATM*-loop ensures that p53 is not decreased due to stress-unrelated or accidental degradations of p53. In a way the ATM*-loop acts as a one-way concentration valve, not allowing that p53 concentrations are decreased below a certain minimum level. The stress-dependent increase of the p53 set-point by the ATM*-p53 loop is a nice example of what Mrosovsky [61] has termed *rheostasis*. Rheostasis is defined as a homeostatic system when the set-point is changed but defended in relationship to a changed external or internal environmental condition or due to stress. A typical example of rheostatic regulation is the defended increase of body temperature (fever) due to an infection. For more examples of rheostasis, see Ref [61].

As indicated earlier [53], the role of the Mdm2-p53 loop seems to avoid premature apoptosis by counteracting uncontrolled rises of p53 above the Mdm2-determined set-point. However, the set-point of the Mdm2-p53 controller does not seem to be fixed. Peng et al. [58] found that treating DLD1 cells with the DNA-damaging agent camptothecin (CPT) led to a decrease in Mdm2 levels. The authors’ interpretation was that Mdm2 degradation under DNA-stress is actually promoted. Such an increase in the Mdm2 degradation (*k*_4_) by DNA-damaging conditions would lead to an increased p53 set-point of the Mdm2 controller (Eq 23), which ultimately could reach apoptotic p53 levels. Chaperones and HSP90 lead to an additional stabilization of p53.

Thus, our model suggests that the individual feedback loops act as temporary stabilizers of p53 when DNA stress is encountered. They result in a gradual step-wise increase in p53 concentration, where each step is under homeostatic (rheostatic) control. When DNA repair is successful and stress levels are removed, p53 concentration falls back to its minimum set-point determined by the Per2-p53 loop. In principle, additional, not yet identified negative feedback loops of p53 with other controllers could be involved in such a rheostatic regulation of p53 during DNA stress. Considering the “plethora of proposed feedback interactions” of p53 [64], an investigation of additional feedbacks in terms of their inflow/outflow behavior may provide further insights and novel suggestions about the workings between different controllers in the p53 network.

## Why oscillations?

The here presented homeostatic (rheostatic) model on how p53 levels are controlled does not necessarily need to involve oscillations. As shown in Fig 4 both the oscillatory and the non-oscillatory versions of the coupled homeostats work equally well. The same goes for the p53 system (Fig 8) when the ATM* and Mdm2 controllers are in a non-oscillatory mode (S3 Text). Thus, sustained or damped oscillations could simply be a byproduct of the negative feedback loops. What supports partly such a view is that a large fraction of *γ*-irradiated cells (≈ 40%) do not show oscillations [52] and that there is a considerable heterogeneity of p53 dynamics even in genetically identical cells [64]. p53 in nonoscillatory cells may still be regulated by the same feedback loops as in oscillatory cells with the same regulatory outcome (cell death or cell survival). The large heterogeneity of p53 dynamics seems to indicate that there is no selective advantage whether homeostatic control of p53 in some instances occurs oscillatory while under other conditions it does not. Porter et al. found that physiologically relevant DNA damage responses apparently begin already after very few p53 pulses or even before the first p53 pulse is completed, and that coordination of p53 target genes increases with successive p53 pulses [65]. This observation can be interpreted in such a way that the coordination of p53 target genes become established when p53 approaches a stress-level dependent steady state/set-point with the suggested [51] possibility that p53 pulses and their dynamics trigger different signal transduction pathways.

There is also the possibility that some controllers (for example Per2 and Mdm2) are oscillatory (due to zero-order degradation with respect to p53), while the ATM* feedback loop is non-oscillatory (due to first-order degradation of p53). This would explain the observation that a certain fraction of cells, by being less susceptible towards gamma irradiation, are controlled by ATM* and thereby are non-oscillatory, while the other part of cells which is more sensitive towards radiation has p53 control by Mdm2 and shows oscillations.

## Summary and conclusion

We have shown that oscillatory homeostats can impose specific frequencies on the oscillations of a controlled variable. In case of perturbations or stress acting on the controlled variable a switch between one oscillatory controller to another is accompanied by a corresponding switch in frequency in the controlling feedback. By analyzing the inflow/outflow control structures of three p53 negative feedback loops (Per2, ATM*, and Mdm2) we were able to assign certain functionalities to each of them. Per2 provides circadian inflow control over p53 by keeping it at the lowest set-point level, ensuring that p53 can be rapidly up-regulated in case of DNA damage/stress. In case of DNA damage the ATM*-p53 feedback loop, another inflow controller, leads to increased p53 levels depending on the stress level. Since the ATM*-induced p53 concentrations are under homeostatic control and defended, the ATM*-p53 feedback loop provides a nice example of what Mrosovsky has termed rheostasis. As an outflow controller, the Mdm2-p53 loop does not allow that p53 levels are raised above the controller’s set-point, probably to avoid premature apoptosis. However, additional mechanisms, such as chaperones and heat shock proteins, in particular HSP90, seem to increase the controller’s set-point and stabilize p53, which finally may lead to apoptosis.

We have considered here only negative feedback loops without the addition of feedforward or positive feedback. While the circadian rhythms show limit-cycle behavior, the p53/ATM* and p53/Mdm2 oscillations are conservative. Including positive feedbacks into the model will certainly make changes in the network’s dynamic and may lead to limit cycle behavior where conservative oscillations are observed. However, we do not think that the qualitative (homeostatic) properties of the interacting negative feedback loops will be significantly altered. Although the p53 model presented here is a far cry from what goes on in a real cell, the inflow/outflow approach and the conditions how negative feedback loops can interact without dysfunctional behavior (wind-up) appears to be an alternative and novel aspect how t analyze the organization of feedback loops in cells and organisms.

## Supporting information

**S1 Matlab. Matlab programs.** A zip-file with Matlab programs showing the results from Figs 4a-b, Figs 9a-d, and Figs 10a-c (left panels).

**S1 Gnuplot. Interactive visualization.** A zip-file with a gnuplot script and an avi video file showing an interactive view of Fig 3.

**S2 Gnuplot. Interactive visualization.** A zip-file with a gnuplot script and avi video files showing interactive views of panels a-d in Fig 5.

**S1 Text. Determination of set-point and period length of the ATM* controller.**

**S2 Text. Combined m1 and m5 controllers and their cooperative and wind-up behaviors.**

**S3 Text. Steady state levels of model (Fig 7) as a function of stress parameter *k*_30_ when feedback loops are non-oscillatory.**

